# Temporal population structure of invasive Group B *Streptococcus* during a period of rising disease incidence shows expansion of a CC17 clone

**DOI:** 10.1101/447037

**Authors:** Dorota Jamrozy, Marcus C de Goffau, Merijn W Bijlsma, Diederik van de Beek, Taco W. Kuijpers, Julian Parkhill, Arie van der Ende, Stephen D. Bentley

## Abstract

Group B *Streptococcus* (GBS) is a major cause of neonatal invasive disease worldwide. In the Netherlands, the incidence of the disease increased, despite the introduction of prevention guidelines in 1999. This was accompanied by changes in pathogen genotype distribution, with a significant increase in the prevalence of isolates belonging to clonal complex (CC) 17. To better understand the mechanisms of temporal changes in the epidemiology of GBS genotypes that correlated with the rise in disease incidence, we applied whole genome sequencing (WGS) to study a national collection of invasive GBS isolates. A total of 1345 isolates from patients aged 0 – 89 days and collected between 1987 and 2016 in the Netherlands were sequenced and characterised. The GBS population contained 5 major lineages representing CC17 (39%), CC19 (25%), CC23 (18%), CC10 (9%), and CC1 (7%). There was a significant rise in the prevalence of isolates representing CC17 and CC23 among cases of early-and late-onset disease, due to expansion of discrete sub-lineages. The most prominent was shown by a CC17 sub-lineage, identified here as CC17-1A, which experienced a major clonal expansion at the end of the 1990s. The CC17-1A expansion correlated with the emergence of a novel phage carrying a gene encoding a putative adhesion protein, named here StrP. The first occurrence of this phage (designated phiStag1) within the collection in 1997, was followed by multiple, independent acquisitions by CC17 and parallel clonal expansions of CC17-1A and another cluster, CC17-1B. The CC17-1A clone was identified in external datasets, and represents a globally distributed invasive sub-lineage of CC17. Our work describes how a sudden change in the epidemiology of specific GBS sub-lineages, in particular CC17-1A, correlates with the rise in the disease incidence, and indicates a putative key role of a novel phage in driving the expansion of this CC17 clone.

**Author summary:** Group B *Streptococcus* (GBS) is a commensal organism of the gastrointestinal and genitourinary tracts. However, it is also an opportunistic pathogen and a major cause of neonatal invasive disease, which can be classified into early-onset (0 – 6 days of life) or late-onset (7 – 89 days of life). Current disease prevention strategy involves intrapartum antibiotic prophylaxis (IAP), which aims to prevent the transmission of GBS from mother to baby during labour. Many developed countries adapted national IAP guidelines. In the Netherlands, these were introduced in 1999. However, the incidence of GBS disease increased after IAP introduction. In this study we applied whole genome sequencing to characterise a nationwide collection of invasive GBS from cases of neonatal disease that occurred between 1987 and 2016. Analysis of GBS population structure involving phylogenetic partitioning of individual lineages revealed that the rise in disease incidence involved the expansion of specific clusters from two major GBS lineages, CC17 and CC23. Our study provides new insights into the recent evolution of the ‘hypervirulent’ CC17 and describes a rapid expansion of a discrete, pre-existing sub-lineage that occurred after acquisition of a novel phage carrying a putative adhesion protein gene, underscoring the major role of CC17 in neonatal diseases.

## Introduction

Group B *Streptococcus* (GBS) is a member of the human gastrointestinal and genital microbiota, and was first recognised as a pathogen in association with bovine mastitis [1]. Human infections with GBS were only sporadically reported until the 1960s when it emerged as a major cause of neonatal invasive infections in developed countries [2]. Neonatal infections are categorised as either early onset disease (EOD, developed during days 0-6 of life) or late onset disease (LOD, developed during days 7-89 of life). While the EOD is due to intrapartum transmission of GBS from a colonized mother, the routes of GBS acquisition in LOD are poorly understood [3]. Current GBS prevention strategies involve intrapartum antibiotic prophylaxis (IAP) for high-risk or GBS-colonized pregnant women, which has been shown to be effective in reducing the rates of EOD [3]. However, following the implementation of national IAP guidelines in high-income countries (HICs) since the late 1990s, variable outcomes were observed. While a number of studies reported a significant drop in the incidence of EOD [4–7], in some HICs the rates of both EOD and LOD had increased [8–10]. This has raised concerns about the efficacy of the local disease prevention guidelines. In some affected countries such as the Netherlands and the UK, IAP is offered based on risk stratification. This is in contrast to a more widely implemented culture-based screening for GBS carriage during pregnancy. Attention has therefore focused on reassessment of local screening methods [8]. Still, other factors such as changes in the host behaviour or medical practice, and improvements in disease detection might have contributed to the rising incidence of the invasive GBS disease [8]. We showed that in the Netherlands it involved a change in GBS genotype distribution, pointing to a possible role of the pathogen itself.

Among the key molecular surveillance targets for invasive GBS is the capsular polysaccharide serotype (CPS), with 10 CPS serotypes (Ia, Ib, II to IX) currently recognised [11]. CPS III represents the dominant GBS serotype amongst neonatal invasive isolates [12, 13]. Of clinical significance in GBS surveillance is also screening for the presence of genetic variants associated with antimicrobial resistance or reduced susceptibility. Penicillin is the drug of choice for IAP and treatment of GBS infections [14]. Reduced susceptibility to beta-lactam antibiotics is rare and can be associated with mutations in the penicillin binding proteins such as *pbp2x* [15]. Prevalence of GBS resistance to erythromycin and clindamycin, alternative therapeutics for individuals with penicillin intolerance, varies by region and is mediated by carriage of horizontally transferred resistance genes [14]. Although not used against GBS infections, human GBS isolates have a high prevalence of tetracycline resistance genes, due to carriage of clonally-distributed conjugative and integrative elements (ICEs)[16].

The molecular epidemiology of GBS is often studied using multi-locus sequence typing (MLST), which characterizes bacterial isolates based on allelic variation in seven house keeping genes. MLST classification revealed clonal complex (CC) 17 as the most common cause of GBS neonatal invasive disease [17]. Due to the overrepresentation of CC17 amongst GBS isolates from invasive disease in newborns, in particular neonatal meningitis, it is recognised as the hypervirulent clone [18, 19]. However, molecular characterisation of bacterial populations using approaches based on selected loci, such as MLST, offers a limited genotypic information and low degree of discrimination between isolates. Instead, whole genome sequence (WGS) analysis provides unparalleled resolution between isolates and can be utilised to study mechanisms of GBS evolution as well as phylogenetic relationship between isolates from various hosts and sources [20–24]. Here we describe WGS analysis of GBS isolates collected between 1987 and 2016 from cases of neonatal invasive disease in the Netherlands, where a rise in the incidence of the disease has been observed despite the introduction of the national disease prevention guidelines in 1999. The aim of this study was to characterise in detail the population structure of invasive GBS from the Netherlands and understand the mechanisms of the previously reported change in GBS genotype prevalence following implementation of the IAP policy [8].

## Results

### Population structure of invasive GBS and temporal epidemiology of sub-lineages

We performed WGS analysis of a Dutch nationwide collection of 1345 GBS isolates from cases of invasive infection in children aged 0 – 89 days identified between 1987 and 2016 (isolate summary in S1 Table). Analysis of GBS population structure using hierBAPS revealed five major clusters, which correlated with the MLST and a phylogeny based on a core gene alignment (S1 Fig). These clusters represented the five major human-associated lineages (98% of all isolates): CC17 (n=526, 39%), CC19 (n=332, 25%), CC23 (n=239,18%), CC10 (n=127, 9%), and CC1 (n=97, 7%). The majority of CC17, CC19 and CC23 of isolates belonged to the founder ST (ST17 [n=493, 94%], ST19 [n=262, 79%] and ST23 [n=205, 86%], respectively) while CC1 and CC10 represented more heterogeneous lineages with a higher diversity of member STs (S2 Fig). The most common capsular serotype, based on a genotypic analysis, was III (n=817, 61%), followed by serotype Ia (n=262, 19%). Other serotypes were represented by up to 6% of isolates, which included Ib (n=84, 6%) II (n=76, 6%), V (n=64, 5%), IV (n=29, 2%), VI (n=5, 0.4%), IX (n=4, 0.3%) and VII (n=2, 0.1%). All but one CC17 isolates carried capsule serotype III (n=525, 99%), which was also observed in the majority of CC19 isolates (n=278, 86%) (S2 Table). All but two CC23 isolates carried serotype Ia (n=237, 99%) (S2 Table). Within each CC, the majority of isolates that shared a ST also carried the same serotype. However, diversity of serotypes within a ST was also observed, in particular among isolates representing ST1, ST2, ST10, ST12 and ST19, with each ST associated with at least three distinct serotypes (S2 Table).

Analysis of antimicrobial resistance gene carriage showed that 85% of all isolates carried at least one tetracycline resistance determinant (*tetM, tetL, tetO* or *tetW*) (S3 Table). Prevalence of *tet* genes varied between CCs and was highest in CC17 (n=504, 96%), followed by CC23 (n=221, 92%), CC19 (n=272, 82%), CC1 (n=72, 74%) and CC10 (n=61, 48%). A total of 8% of all isolates carried at least one macrolide-lincosamide-streptogramin (MLS) resistance gene (*ermB, ermTR, mefA, msrD, lsaC, lsaE* or *lnuB*) (S3 Table). Carriage of MLS resistance genes was highest in CC1 (n=19, 20%), followed by CC19 (n=30, 9%), CC17 (n=43, 8%), CC23 (n=12, 5%) and CC10 (n=5, 4%). Finally, only two isolates had mutations in the *pbp2x* gene previously associated with reduced susceptibility to beta-lactams (T394I and Q557E) [15], each from CC19.

The number of invasive GBS isolates submitted to the reference laboratory increased significantly over time (p < 0.001, Fig 1), reflecting a rise in the incidence of both GBS EOD and LOD disease in the Netherlands during the study period (S3 Fig). This correlated with a significant rise in isolates belonging to CC17 and CC23 (p < 0.001, Fig 1), while the frequency of other CCs remained stable. To determine how the frequency of individual lineages over time correlated with their temporal population structure, a core genome phylogeny of each major CC was reconstructed and partitioned into monophyletic clusters [25]. For CC17, CC19 and CC23, the phylogeny contained a single dominant sub-lineage (Fig 2). The majority of CC17 isolates belonged to sub-lineage CC17-1 (n=427), followed by sub-lineages CC17-2 (n=85) and CC17-3 (n=12) (Fig 2A). Sub-lineage CC17-1 was further partitioned into clusters CC17-1A (n=217) and CC17-1B (n=57). The remaining CC17-1 isolates formed a heterogeneous background population (CC17-1bg, n=153). A significant increase in frequency was observed for CC17-1A and CC17-1B isolates (p < 0.001), with expansion of CC17-1A largely responsible for the overall rise in the frequency of CC17 isolates (Fig 2B). CC19 clustered into five sub-lineages: CC19-1 (n=183), CC19-2 (n=94), CC19-3 (n=20), CC19-4 (n=18) and CC19-5 (n=17) (Fig 2C). Sub-lineage CC19-1 included two distinct clusters, CC19-1A (n=137) and CC19-1B (n=38). There was a significant drop in the frequency of CC19-1A isolates (p < 0.001) although this was offset by a minor rise in CC19-1B (p = 0.03) and CC19-2 (p = 0.04) isolates, the latter becoming the dominant CC19 sub-lineage from 2010 onwards (Fig 2D). CC23 formed two sub-lineages: CC23-1 (n=177) and CC23-2 (n=61), with CC23-1 further sub-divided into CC23-1A and CC23-1bg (Fig 2E). Change in CC23 frequency was associated with a significant rise in the number of CC23-1A isolates (p < 0.001, Fig 2F). The clustering of CC1 and CC10 phylogenies correlated with grouping based on ST assignment, with each lineage clustering into five and six distinct sub lineages, respectively (Fig 2G and 2I). No significant changes in the number of isolates from the different CC1 and CC10 sub-lineages was observed except for a minor increase in the frequency of CC1-1 isolates (p = 0.002, Fig 2H and 2J). Across this invasive GBS population, the most significant change in cluster frequency was displayed by CC17-1A, followed by CC23-1A. Isolates representing cluster CC17-1A were detected infrequently before 1999 but from 2003 it became intermittently the dominant GBS sub-lineage, with 87% of all CC17-1A isolates identified between 2003 and 2016. During that period, CC17-1A was sporadically outcompeted only by CC23-1A.

**Fig 1.**
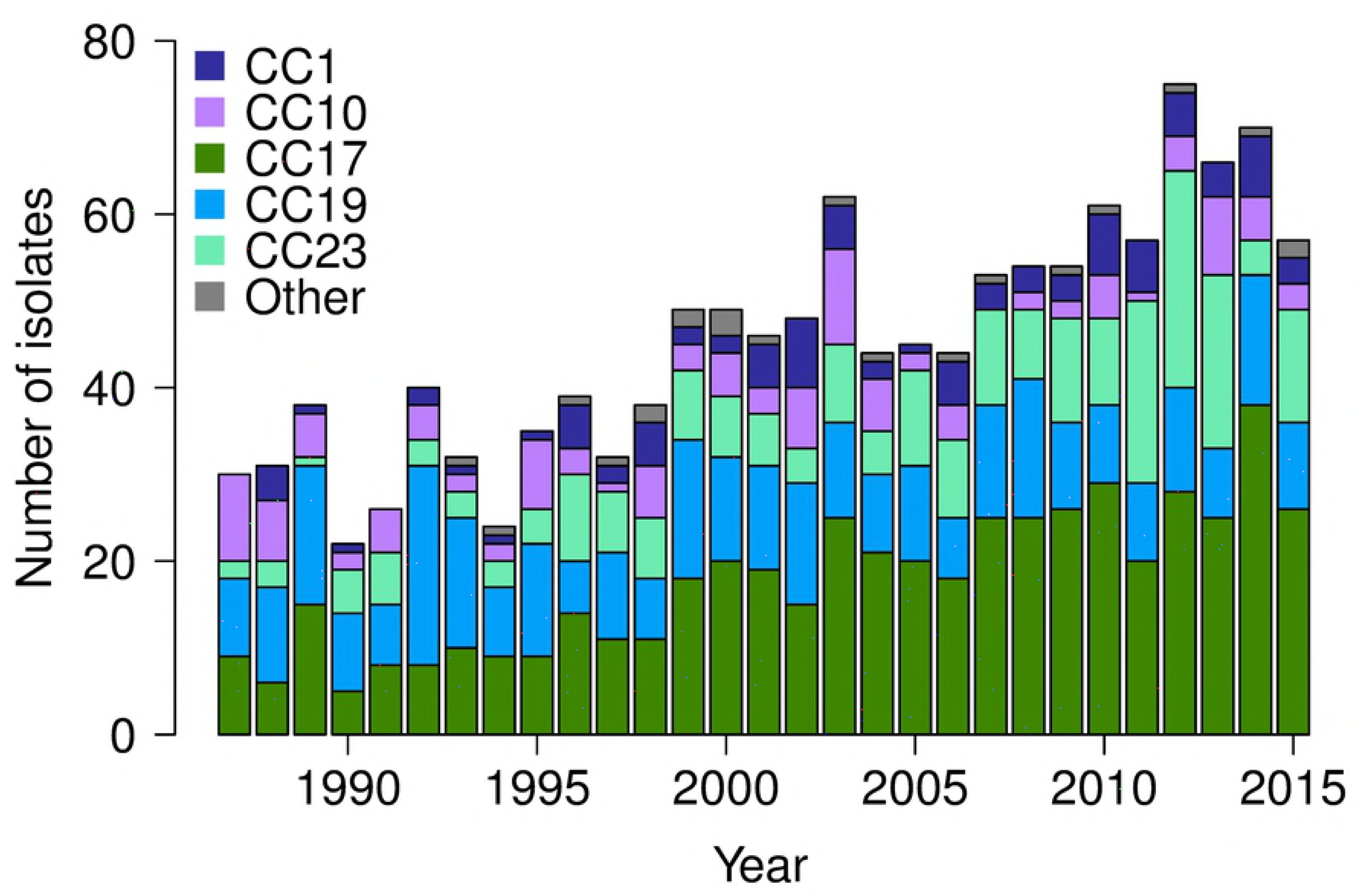
Number of isolates over time stratified by CC. The time scale excludes isolates submitted in 2016 as the sampling period for this study ended in May of that year.

**Fig 2.**
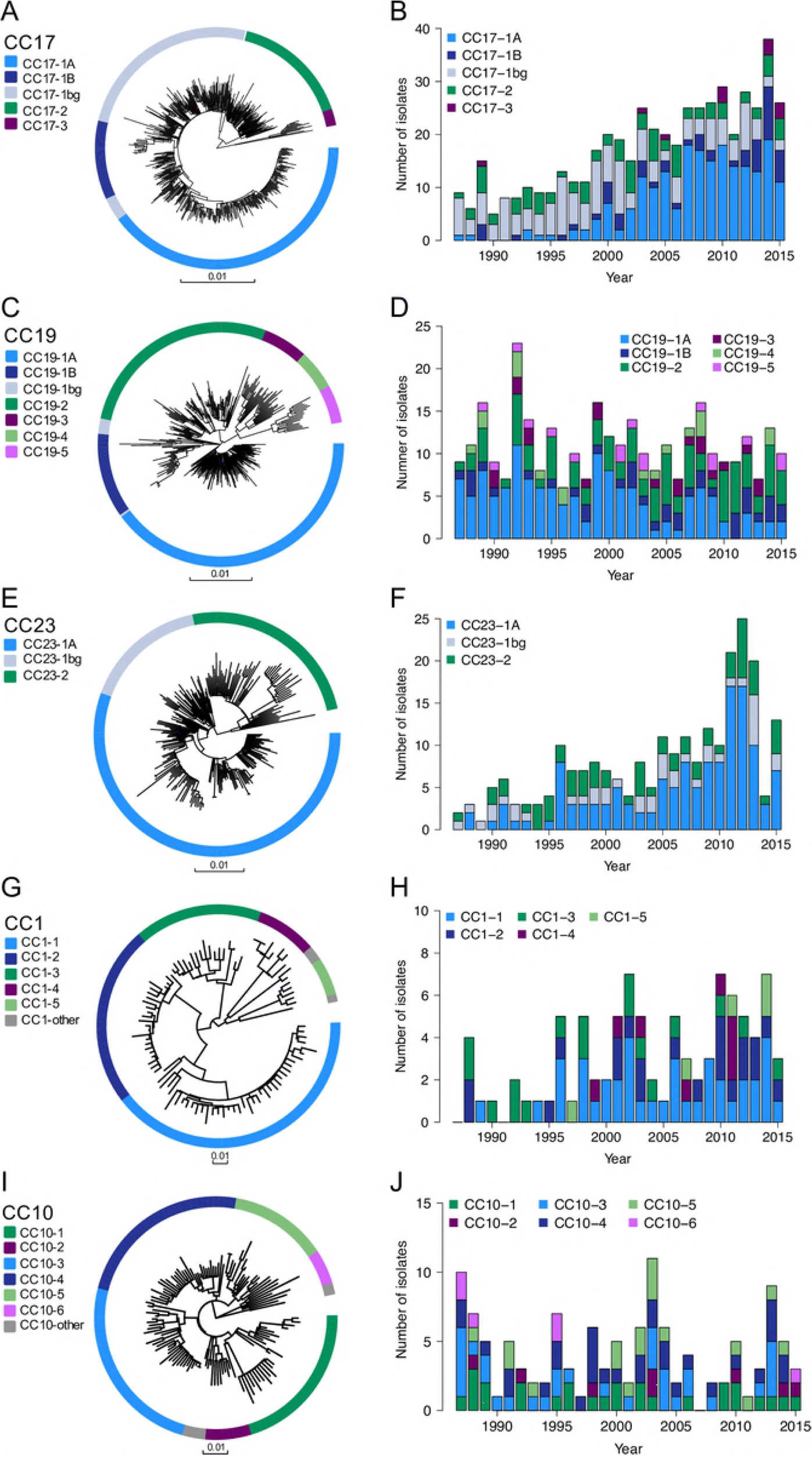
Population structure of GBS CCs and frequency of distinct sub-lineages each year. Core genome phylogenetic tree and the sub-lineage-specific number of isolates each year are shown for CC17 (A, B), CC19 (C, D), CC23 (E, F), CC1 (G, H) and CC10 (I, J). The time scales exclude isolates submitted in 2016 as the sampling period for this study ended in May of that year. Tips of each phylogenetic tree are annotated based on sub-lineage ID (outer ring). Scale bars below each tree represent the number of nucleotide substitutions per site.

Across the collection more isolates derived from EOD (N = 830) than LOD (N = 515). The highest prevalence of EOD isolates was observed for CC1 (81%), followed by CC10 (76%), CC23 (72%) and CC19 (65%) (S4 Fig). CC17 had a significantly lower prevalence of EOD isolates (47%) in comparison with all other CCs (p<0.001). Within each CC, we observed no significant variation in the prevalence of isolates from EOD or LOD between sub-lineages, except for CC23-2 isolates, which were significantly less prevalent amongst LOD cases in comparison with CC23-1A (p < 0.001). There was a significant rise in the number of CC17 isolates among cases of both EOD and LOD over time (p < 0.001) (S5 Fig). For CC23, the rise in the number of isolates was more significant among cases of EOD (p < 0.001) than LOD (p = 0.006) (S5 Fig). We also observed a minor rise in the number of CC1 isolates among EOD cases (p = 0.008) (S5 Fig). For CC17 these temporal changes correlated with the significant increase in the number of CC17-1A isolates amongst cases of EOD as well as LOD (p < 0.001; S6 Fig). For CC23, the rise in CC23-1A isolates was more significant amongst EOD (p < 0.001) than LOD cases (p = 0.04; S6 Fig).

### Evolutionary history of CC17 and CC23

To investigate more closely the association between the rising frequency of CC17 and CC23 among invasive GBS isolates and recent changes in the population structure, a time-calibrated phylogeny and past population dynamics were reconstructed for each lineage based on the Bayesian coalescent skyline model (Fig 3). The estimated mutation rates were 8.51 × 10^-7^ (95% CI 8.1 × 10^-7^ – 8.92 × 10^-7^) and 6.21 × 10^-7^ (95% CI 5.69 × 10^-7^ – 6.73 × 10^-7^) substitutions per nucleotide site per year for CC17 and CC23, respectively. These molecular clock rates correspond to 0.85 and 0.62 SNPs per Mb per year, respectively. Both lineages experienced a significant expansion in the 1960s (Fig 3), which is supported by the epidemiological history of GBS, and is in line with a previous study describing the emergence of adapted GBS clones in mid-20^th^ century [16]. This parallel clade emergence involved multiple sub-lineages within each phylogeny, including CC17-1A (time of the most recent common ancestor, TMRCA: 1968; 95% CI 1965-1970) and CC23-1A (TMRCA: 1968; 95% CI 1965-1972), that have recently increased in prevalence. Both CC17 and CC23 experienced further expansion events in the 1970s and the 1980s. CC23 experienced the most recent clonal expansion towards the end of the 1980s whereas CC17 at the end of the 1990s. The Bayesian coalescent skyline model estimates temporal changes in the effective population size by quantifying the number of coalescent events at discrete time-points. To verify what sub-lineages contributed to specific expansion events in each CC, the frequency of coalescent events over time was compared between the different sub-lineages. This showed sub-lineage-specific peaks that aligned with the expansion events at corresponding time-points (Fig 3). In CC17, the most recent increase in the 1990s was predominantly due to the expansion of CC17-1A, which mirrors the recent rise in frequency of this sub-lineage. For CC23, the most prominent correlation was between the 1980s expansion and the CC23-1A sub-lineage. Annotating tips of CC17 and CC23 phylogenies with the estimated time of tip divergence further demonstrated that the clonal expansion of CC17-1A and CC23-1A occurred across the entire cluster for both sub-lineages (Fig 3).

**Fig 3.**
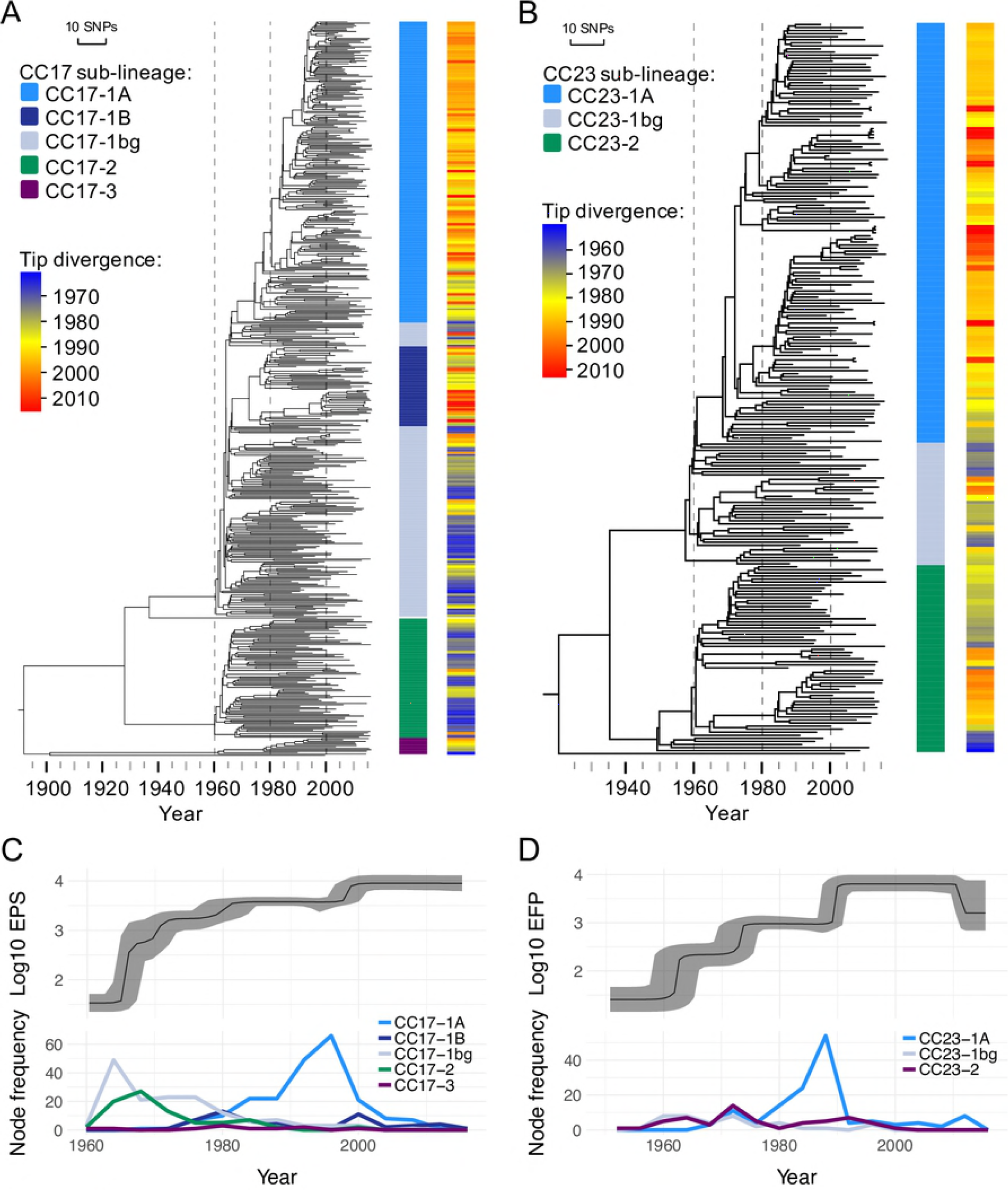
Time-calibrated phylogenies and demographic history of GBS CC17 and CC23. Tips of the maximum clade credibility trees of CC17 (A) and CC23 (B) are annotated with colour strips representing sub-lineage ID, followed by heatmap strips representing the time of tip divergence. Below, the Bayesian coalescent skyline plots show changes (median estimates with high posterior density [HPD] intervals) in the effective population size (EPS) of CC17 (C) and CC23 (D), accompanied by plots of sub-lineage-specific node frequency over time to indicate contribution of distinct sub-lineages to the estimated changes in population size.

### Genomic variation associated with the expanding CC17-1A sub-lineage

To investigate what genomic variants distinguish the CC17-1A sub-lineage, core and accessory genomes of CC17-1A isolates were compared against the background CC17 population. Various single nucleotide polymorphisms (SNPs), occurring on the root of the sub-lineage and the descending branch, separated CC17-1A from the rest of the CC17 population (S7 Fig, S4 Table). Amongst the variants, of particular interest is a double non-synonymous mutation in the *ciaH* sensor histidine kinase gene (position in the COH1 reference genome [accession: HG939456.1]: 954832 and 954833). It occurred on the root of the cluster, and is present in all CC17-1A isolates. Interestingly, the same mutation emerged at least 9 times elsewhere in the CC17 phylogeny, representing a convergent evolution event that might reflect a positive selective pressure acting on this locus (S7 Fig). However, fixation of this variant through clonal expansion was observed only in CC17-1A population. Analysis of association between CC17-1A isolates and variants grouped by occurrence within predicted genes did not identify any specific coding sequences under selective pressure within the CC17-1A cluster. Analysis of loci containing high density of SNPs also did not reveal any regions of homologous recombination associated specifically with the CC17-1A sub-lineage. Finally, we analysed the pan-genome of CC17 to check for variation in the distribution of the accessory genes between CC17-1A isolates and the rest of the CC17 population by performing a principal component analysis (PCA) on the entire accessory gene content of CC17. PCA showed that most of the accessory gene variation within CC17 could be ascribed to the difference between the CC17-1 and CC17-2 sub-lineages (91% of variance explained by PC1). Within the CC17-1 an additional subdivision was visualized by plotting PC2 against PC3 (Fig 4). Three major PCA clusters were composed of isolates from different CC17-1 phylogenetic groups (CC17-1A, CC17-1B and CC17-1bg) indicating that this subdivision was not due to accessory genes unique to the CC17-1A sub-lineage. Correlation analyses of the accessory gene presence/absence within these clusters of isolates revealed the exact combinations of genes responsible for these subdivisions. Specifically, two sets of genes were found to be responsible for the clustering pattern shown in Fig 4. The genes were mapped back on to the CC17 genome assemblies, which showed association with two mobile genetic elements (MGEs). One was an integrative conjugative element, also found in the COH1 reference genome (position: 1847301-1889723) at the 3’ end of tRNA^Lys^ and described previously as ICE_COH1_tRNA^Lys^ [26] (hereafter referred to as ICE_COH1). This MGE encodes several virulence factors such as antigen I/II [27], CAMP factor II [28], and manganese transporter MntH [29]. The second element was a phage, named here phiStag1, also found in the NGBS128 reference genome (accession: CP012480.1, position: 689741 – 748064). It carries a putative adhesion protein gene, named here *strP* (NGBS128 locus tag: AMM49_03725). The protein sequence of StrP contains a ConA-like lectin domain and a C-terminal LPXTG region.

**Fig 4.**
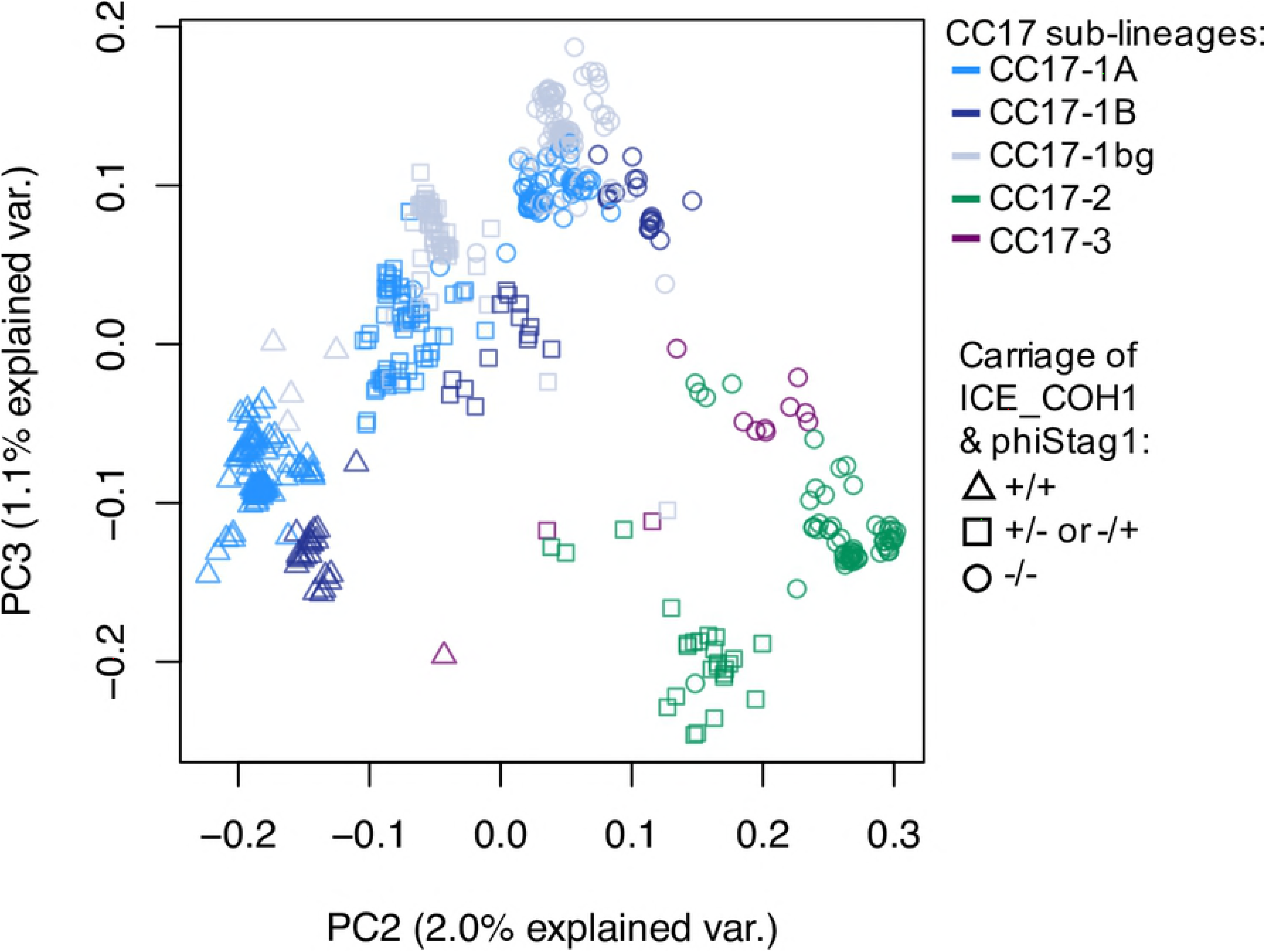
Principal component analysis of the accessory gene distribution in CC17. Each data point is colour-coded to show isolate sub-lineage ID (CC17-1A, CC17-1B, CC17-1bg, CC17-2 or CC17-3). Data point shapes represent the pattern of ICE_COH1 and phiStag carriage to indicate if both MGEs are present (+/+), if only either one MGE is present (+/-or -/+), or if neither MGE is present (-/-). Isolates from CC17-1 sub-lineage revealed all three patterns, although no isolates from CC17-2 carried both MGEs.

A total of 238 and 179 CC17 isolates carried ICE_COH1 and phiStag1, respectively. The composition of PCA clusters reflected the distribution of these MGEs across the CC17 population, with isolates grouping together based on carriage of both, either one or none of the MGEs (Fig 4). MGEs were intermittently distributed across the CC17 phylogeny, demonstrating multiple, independent horizontal transfers (S8 Fig). However, clonal expansion by phiStag1-positive isolates was observed only in clusters CC17-1A and CC17-1B (S8 Fig), both showing a significantly higher prevalence of phiStag1 than either CC17-1bg or CC17-2 (p<0.001). In contrast, clusters of ICE_COH1-positive isolates occurred across the CC17-1 sub-lineage and were not unique to either CC17-1A or CC17-1B (S8 Fig). Instances of MGE co-carriage were mostly sub-lineage-specific as presence of both MGEs was significantly more common than carriage of either MGE in CC17-1A (OR=63, p<0.001) and CC17-1B (OR=87, p<0.001). Analysis of MGE carriage over time revealed that the ICE_COH1 element was present in CC17 isolates throughout the sampling period (S9 Fig). In contrast, phiStag1 first occurred in the analysed CC17 population in 1997. The emergence of phiStag1 in CC17 coincides therefore with the onset of CC17-1A clonal expansion, with the frequency of phiStag1 increasing rapidly in CC17 after 2000 (S9 Fig).

### Epidemiology of the phiStagl phage and *strP* gene

To understand better the origins of the phiStag1 phage, we screened other isolates within our collection, and found that it occurred sporadically in other CCs, most frequently in CC23 (n=41, 17%). The first phiStag1 positive isolate in our GBS collection was CC17 collected in 1997, strengthening the observation that phiStag1 emerged recently in this GBS population. We further expanded the analysis dataset by incorporating published genomic data of 4112 globally distributed GBS isolates from human and animal sources (S5 Table). The phiStag1 phage occurred most commonly in CC17 isolates (25%), followed by CC23 (10%), CC10 (7%), CC19 (6%), and CC1 (4%) (S5 Table).

For CC17, we combined 722 external and 526 Dutch isolates to reconstruct a global phylogeny of CC17. The majority of global isolates co-clustered with the Dutch samples, with 340 and 354 of isolates representing sub-lineages CC17-1 and CC17-2, respectively (S10 Fig). We found that the occurrence of phiStag! phage among the global CC17 isolates was associated predominantly with CC17-1 isolates, more specifically sub-lineages CC17-1A (67% of all CC17-1A isolates) and CC17-1B (74% of all CC17-1B isolates) (S5 Table). Analysis of the global collection also revealed a 95% prevalence of the *strP* gene among bovine-associated isolates from various CCs (S5 Table). However, the gene occurred on a phage that shared less than 90% sequence identity with phiStag1. A historical bovine isolate from 1987 was the earliest *strP*-positive sample identified in this global collection. To investigate further the occurrence of *strP* in historical GBS samples, we screened published reference genomes from the National Collection of Type Cultures (NCTC), available through the NCTC 3000 Project (https://www.phe-culturecollections.org.uk/collections/nctc-3000-project.aspx). The *strP* gene was identified in GBS samples collected before 1950, including strains NCTC 6175, NCTC 8100, and NCTC 8183, all collected from either bovine or unknown host. This suggests a long-term association of the *strP* gene with the bovine GBS population, before its dissemination into the human-associated GBS via a phage.

## Discussion

WGS analysis of a national collection of invasive GBS from cases of neonatal infections that occurred over the last three decades in the Netherlands revealed clonal expansion of discrete sub-lineages from CC17 and CC23. These specific sub-lineages, CC17-1A and CC23-1A, did not represent recently diverged clones as time-calibrated phylogenies estimated the TMRCA for each cluster around the second half of the 1960s. The significant increase in prevalence of CC17 and CC23 during the study period was therefore associated with a recent expansion of clones that have existed since GBS first emerged as a major neonatal pathogen. A sudden, parallel increase in the prevalence of specific clones might result from changes in environmental pressures such as antibiotic usage, which would select for a pre-existing, better adapted clone [30]. Clonal expansions of CC17 and CC23 occurred following the introduction of national GBS disease prevention guidelines in the Netherlands in 1999. However, a link between the IAP and the expansion of these GBS sub-lineages could not be established here as no data was available on whether mothers received the IAP and so it was not possible to determine which EOD cases represented an IAP failure rather than absence of IAP treatment. In cases associated with the latter, no antimicrobial exposure would have occurred unless other antimicrobial treatment was received. Finally, analysis of genomic variants associated with CC17-1A did not reveal any significant association with antimicrobial resistance genes or resistance-conferring SNPs.

Sudden clonal expansion of a bacterial pathogen might be also mediated by genetic changes within circulating clones involving acquisition of novel genes [31]. An increase in prevalence of a pre-existing GBS clone after it acquired new genetic determinants was reported recently for GBS ST-1 serotype V, associated with invasive disease in non-pregnant adults [32, 33]. Our work demonstrated that the expansion of CC17-1A clone, which became the dominant GBS sub-lineage during the 2000s, correlated with the acquisition of the phiStag1 phage. This scenario is further supported by association of phiStag1 with another expanding cluster, CC17-1B. Although the exact role of phiStag1 is unclear, it carries a putative adhesion protein gene, *strP*. Acquisition of a gene encoding a novel cell-surface protein was also linked with the expansion of the GBS ST-1 serotype V [33], which together with our data points to a potentially key role the transfer of such genetic variants might play in the evolution of GBS virulence. This is not unprecedented, as acquisition of phage-associated virulence factors has been linked with the re-emergence and persistence of a hyper-virulent *S. pyogenes* clone [31]. However, further work to describe the function of StrP is required. Presence of the C-terminal LPXTG motif characterises cell wall-anchored proteins whereas the N-terminal ConA-like lectin domain can be found in staphylococcal adhesion proteins such as the platelet-binding SraP from *S. aureus* [34]. The high prevalence of a homologous phage with the *strP* gene in bovine-associated GBS isolates might also hint at its possible biological function. In a bovine host, GBS is an obligate udder pathogen causing persistent mastitis [35]. Virulence of GBS is dependent on ability to adhere to the bovine mammary tissue [35], which in turn requires specific adhesion determinants.

No GBS vaccine is currently licensed, however, maternal vaccination may be a better alternative to IAP for prevention of neonatal GBS infections. Vaccination implementation is likely to have a critical impact on GBS clone distribution, which might involve selection of non-vaccine variants. Better understanding of GBS population structure equips us with a genomic framework for future studies, in particular monitoring shifts in clonal distribution following vaccine introduction. Particular focus on the epidemiology of the expanding CC17-1A sub-lineage is also warranted. We identified the presence of this clone in other GBS collections including a high prevalence amongst invasive isolates from China and Canada [20, 36]. Further international genomic GBS surveillance data is needed to better understand the contribution of CC17-1A sub-lineage to the burden of GBS invasive disease in newborns outside of the Netherlands.

## Materials and methods

### Bacterial isolates

The isolate sampling framework for this study has been described previously [5]. Isolates were derived from a nationwide surveillance study of bacterial meningitis and infant bacteraemia conducted by the Netherlands Reference Laboratory for Bacterial Meningitis. A total of 1345 samples, isolated between 24.01.1987 and 25.05.2016, from patients aged 0 – 89 days were included in the study. Isolates were derived from invasive infections, majority defined as a positive culture from blood or cerebrospinal fluid, with three samples isolated from other body sites (S1 Table). Patient age was calculated as the number of days between date of birth and the earliest known date of the illness; mostly the first date of culture. Early-onset disease (EOD) was defined as invasive infection between 0 – 6 days after birth whereas late-onset infection was defined as invasive infection between 7 and 89 days of life.

### Additional genome sequence data

To complement the analysis, published whole genome sequence data of 4112 GBS isolates from previous studies was also included in this work (S5 Table). This was composed of isolates described by Da Cunha et al. (n=228) [16], Rosini et al. (n=127) [37], Flores et al (n=184) [33], Seale et al (n=1034) [38], Teatero et al (n=77) [20], Campisi et al (n=84) [22, 36], Almeida et al. (n=192) [21, 23], Metcalf et al. (n=2025) [15], and Kalimuddin et al (n=161) [24].

### Whole genome sequencing

Genomic DNA was extracted using Wizard^®^ Genomic DNA Purification Kit (Promega). Tagged DNA libraries were created according to the Illumina protocol. Whole-genome sequencing was performed on the Illumina HiSeq 2000 platform with 125 bp paired-end reads. Annotated assemblies were produced using a pipeline described previously [39]. For each sample, sequence reads were used to create multiple assemblies using VelvetOptimiser v2.2.5 (https://github.com/tseemann/VelvetOptimiser) and Velvet v1.2 [40]. An assembly improvement step was applied to the assembly with the best N50 and contigs were scaffolded using SSPACE [41] and sequence gaps filled using GapFiller [42]. Automated annotation was performed using PROKKA v1.11 [43] and a *Streptococcus*-specific database from RefSeq [44].

### Isolate genotype analysis

GBS population structure was determined with hierBAPS [45] based on the first level of BAPS clustering. Multi-locus sequence typing, based on the *S. agalactiae* MLST scheme [17], was performed using SRST2 [46]. Isolates were assigned to a clonal complex if they represented the same hierBAPS cluster as the founder ST. The founder ST was confirmed with eBURSTv3 [47]. To verify that isolates sharing CC assignment belonged to the same monophyletic group, a core gene alignment was generated using Roary [48] and an approximately maximum-likelihood tree was reconstructed using FastTree [49]. The tree was visually inspected to confirm that isolates sharing CC assignment belonged to the same clade. Isolates that shared the same hierBAPS cluster but were phylogenetically distinct were assigned to a unique CC based on their ST.

Isolate CPS serotype was determined by *in silico* PCR using previously described primers for detection of serotypes Ia, Ib and II-IX [50, 51]. Carriage of antimicrobial resistance genes was checked with SRST2 [46] using the ARG-ANNOT database [52], supplemented with sequences of GBS resistance determinants described by Metcalf et al [15].

### Phylogenetic tree reconstruction

To reconstruct phylogeny of CC1, CC10, CC17, CC19 and CC23, sequence reads were mapped using SMALT v0.7.4 (https://www.sanger.ac.uk/science/tools/smalt-0) to CC-specific *S. agalactiae* reference genomes: SS1 (CC1, accession: CP010867), Sag37 (CC10, accession: CP019978), NGBS128 (CC17, accession: CP012480), H002 (CC19, accession: CP011329) and NEM316 (CC23, accession: AL732656), respectively. For each CC, a core genome alignment was created after excluding MGE regions, variable sites associated with recombination (detected with Gubbins [53]) and sites with more than 5% proportion of gaps (i.e. sites with an ambiguous base). A maximum likelihood (ML) phylogenetic tree was generated with RAxML v8.2.8 [54] based on a generalised time reversible (GTR) model with GAMMA method of correction for among site rate variation and 100 bootstrap (BS) replications. Intra-CC phylogenetic clustering was performed on ML phylogenetic trees, which were partitioned using incremental pairwise SNP distance [25]. Cluster reliability was based on a BS support of ≥90.

For time-calibrated phylogenetic tree reconstruction of CC17 and CC23, each sequence in the CC-specific core genome alignment was annotated with the date of isolation based on day, month, and year. A correlation between root-to-tip distance and sampling date was checked using TempEst [55]. Bayesian inference of phylogeny and past population dynamics was performed with BEAST 2 [56] under a GTR+Γ nucleotide substitution model, strict clock with coalescent Bayesian skyline model. The MCMC chain was run for 50 million generations, sampling every 1000 states. Log files from five independent runs were combined after removal of burn-in (5% of samples) using LogCombiner and analysed with Tracer v1.5. Maximum clade credibility (MMC) tree was generated with TreeAnnotator. MCC tree statistics were extracted with BEASTmasteR [57] (http://phylo.wikidot.com/beastmaster). All phylogenetic trees were annotated using Evolview [58, 59].

### Core genome polymorphisms

To analyse core genome SNPs in CC17 isolates, sequence reads were mapped using SMALT v0.7.4 (https://www.sanger.ac.uk/science/tools/smalt-0) to the COH1 reference genome (accession: HG939456), and the core genome alignment was prepared as described for the phylogenetic tree reconstruction. The ancestral single nucleotide polymorphisms were reconstructed along each branch of the ML phylogeny using Sankoff parsimony [60–62].

### Accessory genome analysis

Pan-genome analysis of CC17 was performed with Roary [48]. Distribution of genes was investigated with principal component analysis (PCA), performed in R v3.3.3. Genes that were correlated (based on Spearman’s rank correlation coefficient) with selected PCA clusters were identified and analysed further. To determine their genomic context, genes were mapped back to whole genome assemblies using blastn pairwise sequence alignments [63]. Sequence similarity searches of genomic fragments found to contain the accessory genes of interest were performed using the NCBI BLAST and GenBank nucleotide sequence database [64]. MGE and gene carriage was checked by short read mapping with SRST2 [46] with 90% and 99% sequence coverage cut-off, respectively.

### Statistical analysis

All statistical analyses were performed in R v3.3.3. Correlation between genotype prevalence and year of sampling was measured with Mann-Kendall test (*Kendall* package). All tests excluded isolates submitted in 2016 as the sampling period for this study ended in May of that year. Comparison of isolate distribution by the disease onset between lineages, MGE distribution between CC17 sub-lineages and MGE co-carriage was performed with Fisher’s exact test.

## Acknowledgements

The authors like to thank Mrs Sandra Kit-Bovenkerk for her technical assistance, Dr Simon R. Harris and the Sequencing and Pathogen Informatics groups at the Wellcome Sanger Institute for their bioinformatics support.

## Supporting information

**S1. Fig. Population structure of neonatal invasive GBS.**

An approximately maximum-likelihood phylogenetic tree of all GBS isolates based on a core gene alignment, annotated with CC assignment as determined with MLST, hierBAPS cluster ID and CPS serotype. Scale bar below the tree represents the number of nucleotide substitutions per site.

**S2 Fig. ST composition of major CCs.**

Diversity and proportion of STs within CC1 (A), CC10 (B), CC17 (C), CC19 (D) and CC23 (E).

**S3 Fig. Incidence of GBS invasive disease in the Netherlands between 1987 and 2016.**

The incidence of EOD and LOD is shown as the number of cases per 1000 livebirths.

**S4 Fig. Number of isolates from EOD and LOD by CC.**

**S5 Fig. Number of isolates from EOD and LOD each year stratified by CC.**

(A) Number of EOD isolates over time stratified by 5 major CCs. (B) Number of LOD isolates over time stratified by 5 major CCs. Time scale on each plot excludes isolates submitted in 2016 as the sampling period for this study ended in May of that year.

**S6 Fig. Number of CC17-1A and CC23-1A isolates among EOD and LOD cases each year.**

(A) Number of CC17-1A isolates over time stratified by the disease onset. (B) Number of CC23-1A isolates over time stratified by the disease onset. Time scale on each plot excludes isolates submitted in 2016 as the sampling period for this study ended in May of that year.

**S7 Fig. Phylogenetic tree of CC17 and distribution of SNPs associated with CC17-1A sub-lineage.**

Tips of the tree are annotated with colour strips representing (from the left) the CC17 sub lineage ID followed by columns representing the base variant at 20 SNP sites associated with the CC17-1A sub-lineage, occurring on its root (node 1) and the descending branch (node 2). Each SNP is described in more detail in S4 Table. Each column represents a unique SNP site, at the indicated position in the GBS COH1 reference genome. ‘N’ represents isolates with a missing variant at the corresponding site.

**S8 Fig. Distribution of ICE_COH1 and phiStag1 across the CC17 phylogeny.**

Tips of a maximum-likelihood phylogenetic tree of the Dutch CC17 are annotated with strips representing (from the innermost ring): CC17 sub-lineages, presence of phiStag1, presence of ICE_COH1. Scale bar below the tree represents the number of nucleotide substitutions per site.

**S9 Fig. Number of CC17 isolates carrying ICE_COH1 and phiStag1 each year.**

The time scale excludes isolates submitted in 2016 as the sampling period for this study ended in May of that year.

**S10 Fig. Global phylogeny of CC17 with carriage of ICE_COH1 and phiStag1.**

Maximum-likelihood phylogenetic tree of 526 Dutch and 722 globally derived CC17 isolates. Tips of the tree are annotated with strips representing (from innermost ring): CC17 sub lineage, isolate country of origin, presence of phiStag1, presence of ICE_COH1. Scale bar below the tree represents the number of nucleotide substitutions per site.

**S1 Table. Isolates summary**

**S2 Table. Distribution of CPS serotypes by CC and ST.**

For each serotype, the total number of isolates carrying the serotype from the corresponding ST is provided.

**S3 Table. Distribution of antimicrobial resistance genes by major CC.**

The total number of isolates carrying at least one resistance gene to tetracyclines (TET_total), macrolide-lincosamide-streptogramins (MLS_total) and aminoglucosides (AGLY_total) is provided for each major CC. For each antimicrobial class, the identified resistance genes and the number of isolates carrying each gene are also listed.

**S4 Table. List of SNPs associated with CC17-1A cluster.**

**S5 Table. Summary of external GBS genomes included in this study.**

